# MegaPX: fast and space-efficient peptide assignment method using IBF-based multi-indexing

**DOI:** 10.1101/2025.04.14.648734

**Authors:** Ahmad Lutfi, Tanja Holstein, Sandro Andreotti, Thilo Muth

**Affiliations:** Data Competence Center MF2, Robert Koch Institute, Seestraße 10, 13353, Berlin, Germany; VIB-UGent center for Medical Biotechnology, VIB, Belgium; Department of Mathematics and Computer Science, Institute of Computer Science, Freie Universität Berlin, Takustr. 9, 14195, Berlin, Germany

**Keywords:** *k*-mer indexing, Bloom filter, proteomics, metaproteomics, error-tolerant searching, bioinformatics algorithms, similarity search, data structure, mass spectrometry, alignment-free search, *de novo* sequencing, peptide identification

## Abstract

**Motivation:** A central problem for metaproteomic analysis is the often-unknown taxonomic composition of the analyzed microbiomes. Using a database search, the standard approach requires prior knowledge of which proteins and taxa to include in the protein reference database or to use tailored metagenome-derived databases, which are expensive and error-prone in their generation. A possible strategy to circumvent this database search issue is *de novo* sequencing, where peptide sequences are directly identified from mass spectra. However, these sequences must still be mapped back to potentially extensive databases. Here, alignment-based approaches enable robust and precise results, with the potential drawback of high memory usage and long run times.

**Results:** We present MegaPX, a software for rapidly classifying *de novo* peptide sequences against large protein databases. MegaPX implemented as a C++-based tool, uses an alignment-free, *k*-mer-based approach as a taxonomic classification method with the possibility of generating mutated reference databases for error-tolerant searching. It uses various algorithms, including interleaved Bloom filters, to efficiently compute approximate membership queries, ensuring fast processing times while querying and indexing large databases in a multi-indexing fashion. We demonstrate the potential of MegaPX by analyzing different samples, including metaproteomics, against extensive reference databases, highlighting its use as a fast screening tool.

**Availability and implementation:** MegaPX’s source code and all related documentation files are freely available in a GitHub repository (https://github.com/rki-mf2/MegaPX) under the MIT License.

**Supplementary information:** All supplementary tables and figures are included at the end of the preprint.

## Introduction

Proteomics is a rapidly advancing and powerful approach for studying the entire proteome—the complete set of proteins expressed by an organism, tissue, or cell—or specific subsets thereof. This technique provides a detailed and informative landscape of expressed proteins and their modifications under various conditions [HA14], providing valuable insights into biological processes. Proteomics is the crucial starting point in various biological fields and is utilized in medicine, food microbiology, pharmacy, industry, and drug development [KRR19] . Metaproteomics, which is an extension of proteomics, focuses on the large-scale study of protein expression in microbial communities, enabling the characterization of microbial diversity, metabolic activity, and host-microbe interactions. Unlike traditional proteomics, which targets a single organism, metaproteomics analyzes complex microbial environments, making it a valuable tool in fields such as environmental microbiology, human microbiome research, and infectious disease diagnostics [Kle19; Van+21]. It often includes the quantitative profiling of proteins, interactions, and modifications using mass spectrometry-based analytics [SM20] . Among these, liquid chromatography-tandem mass spectrometry (LC-MS/MS) is a high-performance analytical tool increasingly employed in a variety of research and diagnostic applications [Tho+22]. The most commonly used method in protein analysis is the bottom-up approach [El +18] (Suppl. Mat. Figure S1). This method begins with extracting proteins from the target tissue or cell, followed by enzymatic cleavage using a specific enzyme, typically trypsin, which efficiently and precisely cleaves proteins into shorter peptides. These peptides, usually ranging from 5 to 30 amino acids, are separated by liquid chromatography. In tandem mass spectrometry (MS/MS), peptides are first ionized, generating so-called precursor ions, which are then introduced into the mass spectrometer [Mit15] . The MS1 spectrum records the mass-to-charge ratios (m/z) of these precursor ions, providing insights into their molecular composition. Selected precursor ions are then fragmented, producing MS2 (or MS/MS) spectra. These fragment spectra are analyzed using computational methods, most commonly database searching or *de novo* peptide sequencing, to identify and quantify peptides in the sample [WW13] . This process is invaluable for determining protein functions in cells and tissues [FEB10], the discovery of biomarkers [Kar+20a], and the discovery of drugs [Wan+22]. Search engines that rely on MS/MS spectra and a protein sequence database as input are called database-dependent search engines [Mat13], based on comparing experimental spectra, which contain the masses and abundances of peptide fragments, with the theoretical spectra derived from protein sequence databases [Pev+01] . Over the last decade, numerous tools have been developed to enable fast and memory-efficient identification of proteins from MS data. Among these, algorithms such as Andromeda [Cox+11], MS-GF+ [KP14], X!Tandem [CB03], Mascot [BHC09], and InsPecT [Tan+05] match MS/MS spectra to peptides derived from protein sequence databases. However, these approaches face several challenges. The growing size of protein sequence databases, often containing millions of entries, increases computational demands and impacts statistical validation. Additionally, databases may be incomplete, lacking certain proteins or splice variants. Many peptide sequences are also highly similar due to common sequence motifs, post-translational modifications (PTMs), or housekeeping genes. Furthermore, MS data itself is often noisy, incomplete, or contains overlapping peaks, complicating accurate peptide identification [Ver+16] .

In contrast to database searching, *de novo* peptide sequencing aims to identify peptides directly from MS/MS data without relying on protein sequence databases. Recent advances in machine and deep learning have enabled real-time peptide sequence prediction by training on millions of annotated spectra. This approach is particularly valuable when analyzing samples with unknown or highly variable proteomes, such as cancer cells or viral populations, where frequent mutations can hinder traditional database search methods. However, despite their advantages, *de novo* sequencing methods are limited by the availability of comprehensive annotated MS datasets. The absence of datasets covering all possible mutations and PTMs restricts their ability to fully characterize the diversity of viral and cancer-associated proteomes. Several *de novo* sequencing algorithms have been developed, including pNovo3 [Yan+19], SMSNet [Kar+19], Novor [Rap24], DeepNovo [Tra+19], PointNovo [Qia+21], Casanovo [Yil+24], InstaNovo [Elo+23] and Spectralis [Kla+23] . While these methods excel at identifying peptide sequences from MS/MS spectra, they do not inherently determine the corresponding proteins. As a result, *de novo* sequencing alone is insufficient for full proteome characterization. To bridge this gap, these algorithms must be integrated with sequence analysis methods that map identified peptides to proteins using reference databases.

Numerous bioinformatics tools focus on sequence comparison techniques to identify and quantify samples in databases. Comparing sequences with each other plays a crucial role in understanding cellular processes, gene identification, dataset annotation, and pathogen detection [SU19] . By combining *de novo* sequencing with advanced sequence comparison methods, researchers can enhance the accuracy and depth of proteomic studies, particularly in complex or poorly characterized biological systems. Sequence comparison methods typically rely on alignment-based approaches, which determine similarities by evaluating matches, mismatches, and gap positions. These methods are known as local [Smi81] and global [Nee70] alignment depending on the specific application. While alignment-based sequence comparison is accurate for identification, it suffers from quadratic time complexity, as comparisons are performed at the string level and runtime increases exponentially with the number of compared sequences [Rub+17] .

To address these limitations, alternative approaches based on short, overlapping substrings, called *k*-mers, of length *k* are increasingly employed in bioinformatics for sequence comparison and identification. *k*-mer-based methods are widely used in genomics [Cho+09] metagenomics [She+22], and pathogen detection [Kat+21]. These methods are often coupled with approximate string-matching data structures such as Bloom filters [Blo70], quotient filters [Ben+12], cuckoo filters [Fan+14], *k*-distance trees [BF79], *q*-gram indices [lok05], and other related techniques.

This work introduces *MegaPX*, an innovative tool for peptide assignment that uses the **I**nterleaved **B**loom **F**ilter (IBF) data structure [Dad+18] at its core, offering superior performance compared to traditional methods. This results in an enhanced sequence-based assignment strategy. By constructing a set of IBFs, MegaPX enables highly efficient and memory-friendly querying of extensive protein databases. To demonstrate its performance, we first evaluate MegaPX using a small cancer dataset to assess its accuracy in protein classification. We then extend our analysis to *de novo* peptides from a metaproteome sample by querying an extensive protein database with over 700 million protein sequences. This experiment highlights the feasibility of our tool for execution on standard notebooks with minimal memory consumption. MegaPX’s source code and documentation are available in a GitHub repository (https://github.com/rki-mf2/MegaPX) under the MIT License, and it can be run on Linux or **W**indows **S**ubsystem for **L**inux (WSL) using conda-based installations.

## Materials and methods

### De novo peptide Sequencing

In general, *de novo* peptide sequencing algorithms are designed to directly determine the amino acid sequence of peptides from MS data without referring to a protein sequence database. Many algorithms are based on machine or deep learning methods and attempt to make predictions in real-time with specific parametric configurations. We combined the sequencing results from two different deep learning models: Casanovo [Yil+24] and SMSNet [Kar+19] . Both tools were executed on a Linux machine (Ubuntu 20.04.6 LTS 64-bit) with standard GPU configurations. SMSNet was executed using a previously published *de novo* sequencing pipeline [Bes+22] . Both deep learning models have been trained on high-resolution MS/MS data from the human proteome using the HCD library from MassIVE [Wan+18], which consists of 1,114,503 annotated peptides. The annotated spectra were split into three sets: Training, Validation, and Test, in a ratio of 98:1:1, ensuring no overlap of peptides between the datasets. The models were trained with a precursor tolerance of 10 *ppm* and a fragment mass tolerance of 0.02 *Da*. Carbamidomethylation of cysteine was set as a fixed modification, whereas oxidation of methionine and deamidation of asparagine and glutamine were set as variable modifications. For the analysis, we opted to use the pre-trained Casanovo weights from the original publication and the SMSNet model from [Bes+22] . This decision was based on their distinct training procedures, ensuring broader coverage of different sequencing scenarios in our combined approach. Since each *de novo* algorithm calculates scores differently, we included the sequence score extracted from the corresponding algorithm in the peptide header to facilitate the retrieval of this information if needed. This integration allowed for a more comprehensive analysis and comparison of results from both tools. The generated FASTA files contain three peptide categories: (i) shared peptides identified by both Casanovo and SMSNet, (ii) unique peptides from Casanovo, and (iii) unique peptides from SMSNet. By incorporating both shared and unique sequences, we aim to capture a more comprehensive peptide landscape. To ensure data quality, only peptides with a confidence score of at least 70% are retained, these scores represent peptide-level confidence scores obtained by averaging the amino acid-level confidence scores from the corresponding model predictions across the peptide sequence. However, in cases where mutations are more frequent, a lower cut-off of 50% is applied to account for sequence variability.

### Classification Task

#### Interleaved Bloom filter

The efficient querying of several Bloom filters simultaneously poses a challenge in various applications. Conventional methods require each Bloom filter to be queried individually and the query strings to be assigned to the respective references individually. Dadi *et al*. [Dad+18] proposed a novel data structure, the **I**nterleaved **B**loom **F**ilter (**IBF**), to address this challenge. In contrast to conventional Bloom filters, an IBF is a combined structure consisting of several Bloom filters, all of which have the same size *n* and a set of hash functions *H*. In IBF, each bit of each Bloom filter *b* is interleaved into a uniform bit vector, resulting in a combination of Bloom filters in an interleaved fashion with a total size of *b × n*. Within the IBF structure, each bit is internally replaced by a sub-bit vector of size *b* where the *i*−*th* bit correlates with the Bloom filter of bin *B*_*i*_. The rate of false positives in the IBF is closely related to the size of each Bloom filter and the number of hash functions *H* used for *m* inserted elements. The basic concept of the IBF is that the individual Bloom filters no longer have to be queried sequentially. Instead, the IBF combines all bins in a single data structure, enabling the entire set to be queried simultaneously. The false positive rate of the IBF can be calculated using the formula from Dadi *et al*. [Dad+18] :

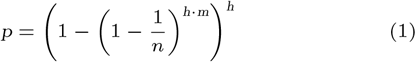

Similarly, the optimal size of each Bloom filter (bin) for a given error rate can be calculated using the equation [Ulr+22] :

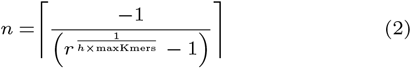

where *r* = 1 − *p*^1*/h*^ and *maxKmers* is the maximum number of *k*-mers in the input user bins. This value is computed by estimating the maximum size of *k*-mers sets in the input user bins. Our tool calculates the size of the IBF based on the vectorized representation of the user bins (input references) using a default false positive rate of 0.01. Figure 1 shows an example of an IBF structure derived from three input reference sequences.

**Fig. 1.**
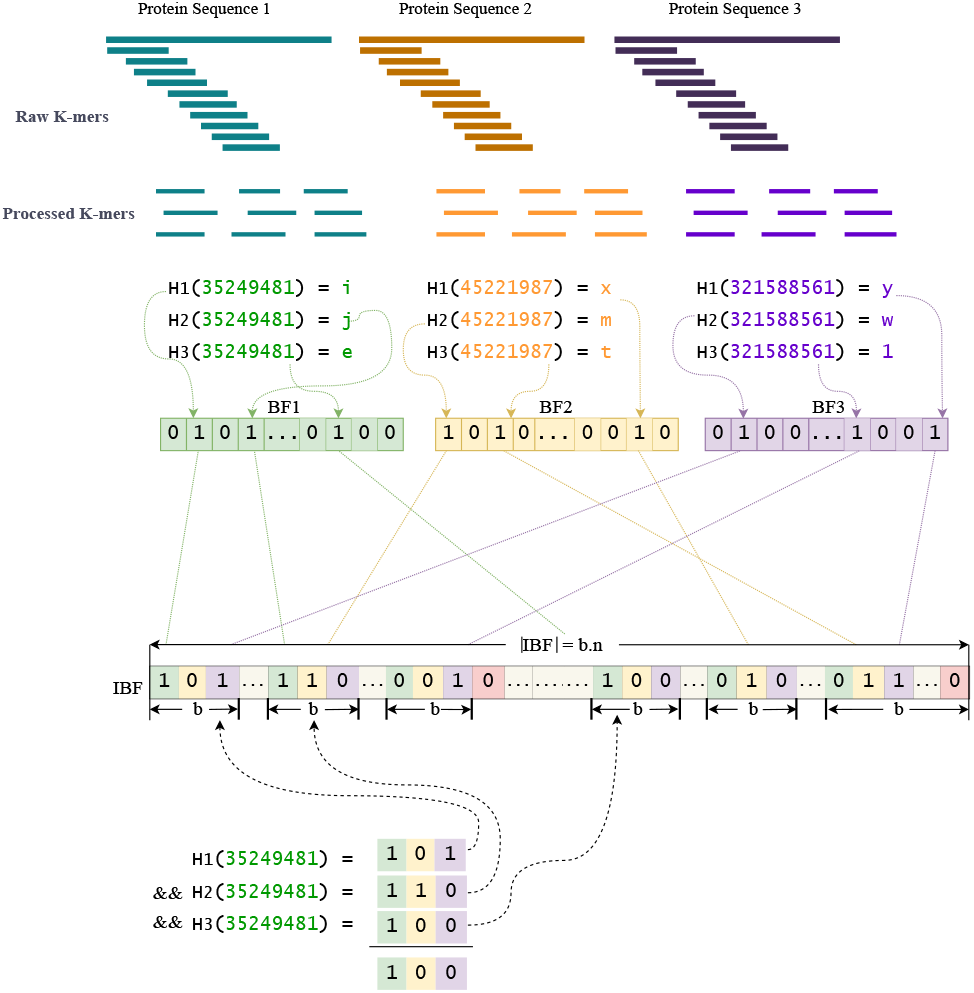
Interleaved Bloom Filter Construction. *k*-mers are generated or processed (mutated) in the first step, depending on the use case. Three positions are then retrieved for each *k*-mer using three different hash functions (*H*_1_ to *H*_3_), with the corresponding bit being set to *1*. These operations are performed independently for each *k*-mer. The resulting Bloom filters, represented by different colors, are then interleaved to construct an interleaved Bloom filter of size *b × n*. When a new reference sequence is inserted into the filter, the size of each sub-vector increases to *b × n*. The sub-bit vectors are combined with a bitwise AND operation for the *k*-mer target or a coded integer *35249481*. This combination occurs after retrieving three positions using the same three hash functions used to construct the IBF, resulting in the required combination vector with the *B*_1_ response.

In the beginning, each reference sequence is *k*-merized to a length *k*. Subsequently, the generated *k*-mers are encoded, and these encoded *k*-mers are either inserted directly into the filter or mutated versions are generated (processed *k*-mers). The resulting *k*-mers are then hashed using three different hash functions, and the resulting hashes of each reference sequence are inserted into the corresponding Bloom filter. These different BFs are then interleaved into a single IBF. However, it should be noted that memory consumption can increase significantly if the reference sequences are of very different sizes, as all BFs are interleaved. Consequently, the size corresponds to the largest BF, meaning other filters also reserve a lot of memory for empty blocks.

#### Peptide classification

Following the query membership of peptide sequences from each sample against a pre-built index, we constructed a count matrix *C*. Each row in this matrix represents a peptide, while each column corresponds to the user bin. The entries of the matrix are the *k*-mers’ hits per bin for every query peptide:

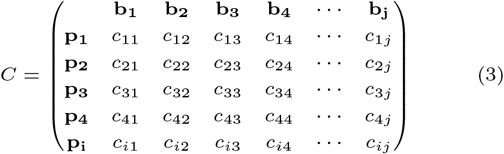

The count matrix is further processed according to the user-defined assignment threshold *D*, representing the percentage of the peptide *P* (number of *k*-mers) that matches the user bin *B* to assign the peptide to that user bin.

Let 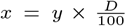, where *y* is the total number of *k*-mers in peptide *P*.

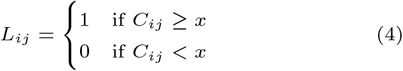

The lookup matrix is built with bits (*1* and *0*) and defined as:

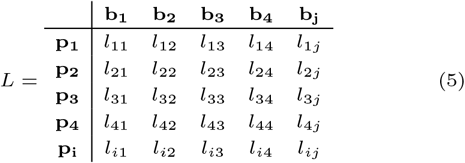

Using the lookup table *L*, the assignment score *S* of the set of peptides in a sample to the corresponding user bin *B* can be computed as:

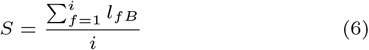

, where *i* is the total number of peptides in the input sample. MegaPX supports querying large databases with errors, enabling an error-tolerant detection of amino acid sequences. Substitution matrices characterize all possible modifications of each amino acid by another and provide essential information for predicting potential changes based on a specific score. The BLOSUM (**BLO**cks **SU**bstitution **M**atrix) series is one of the most widely used substitution matrices. BLOSUM matrices are constructed by clustering aligned sequence segments within conserved blocks based on a specified percentage identity threshold. This process involves analyzing large protein data sets to identify amino acid substitution patterns, which are then used to derive scoring matrices that reflect the likelihood such substitutions occurring in related proteins [HH92]. The scores in BLOSUM matrices are expressed as log-odds values, representing the biological probability of each substitution. These values indicate how frequently a given amino acid replacement occurs in evolutionary-related proteins compared to random chance. BLOSUM62 is derived from sequence blocks with at least 62% identity [HH92]. In a BLOSUM matrix, higher scores indicate a greater probability that the corresponding amino acid substitution occurs during sequence alignment [OCo24].

We generate all possible mutational variants of each *k*-mer by simulating potential amino acid substitutions, guided by sequence similarity, to identify target proteins even in the presence of mutations. The process begins by dividing the target protein sequence into overlapping *k*-mers. For each *k*-mer, we consider all possible amino acid substitutions based on the BLOSUM62 substitution matrix and a user-defined similarity score threshold. Without optimization, this substitution process would require evaluating up to 20^*k*^ combinations, which is computationally intensive and results in long runtimes. To address this, we implemented a pre-sorting step on the substitution matrix. Specifically, for each amino acid, its possible substitution candidates are sorted by their similarity scores in descending order. This sorted matrix enables prioritization of high-scoring substitutions first during the mutational variant generation. In addition, an early stopping mechanism is applied during the substitution process. If a candidate substitution score falls below the user-defined threshold, the loop terminates early, skipping any remaining lower-scoring candidates. This significantly reduces the number of computations required when aligning each mutated *k*-mer to the reference sequence, resulting in a more efficient runtime performance. Furthermore, the generated database contains the original and the mutated *k*-mers, so the original protein sequences are also considered and inserted into the IBF.

#### Multi-indexing classification

MegaPX incorporates a multiple indexing approach to efficiently index and query large databases. In this approach, we partition *n* user-defined references into batches and generate different IBFs, each containing the same number of user bins (determined by the total number of references divided by the split size). Each IBF has its own size depending on the inserted user bins to minimize filter size and reduce corresponding false-positive responses as much as possible. This approach is based on a divide-and-conquer algorithm, where we decompose complex problems into smaller subproblems and combine partial solutions into a comprehensive solution. This strategy aims to reduce the time and memory requirements of processing large data sets. While this approach optimizes indexing efficiency, it may lead to slower query times, particularly when the target database contains a high proportion of over-represented sequences. To address this, MegaPX allows the inclusion of an external blacklist file, enabling users to exclude specific references from the indexing process. These references, often considered outliers, are omitted from subsequent searches to improve query performance.

This method eliminates the need to load, write, and repeatedly query very large IBFs. Instead, it enables querying smaller IBFs, which can be efficiently managed within standard RAM configurations. The workflow of this approach is illustrated in Figure 2 (pseudo-code in Suppl. Mat. Algorithm S1). Additionally, our workflow supports minimizer computation. Minimizers are typically calculated in both the forward and reverse directions of a given sequence, as DNA sequences have palindromic features where the sequence on one strand is the reverse complement of the sequence on the other strand [MS21]. Taking both the forward and reverse strands into account is crucial for applications such as sequence alignment, where similarities can occur on either strand [Min12]. However, unlike DNA sequences, protein sequences are composed of amino acids and do not have a complementary reverse strand. Therefore, when dealing with protein sequences, minimizers are only calculated on the forward strand. In this case, the *k*-mer set is treated as a series of contiguous values, and minimizers are calculated in a single direction. Since protein sequences are generally smaller than DNA or RNA sequences, we have integrated minimizer calculation as an optional feature rather than part of the standard processing pipeline.

**Fig. 2.**
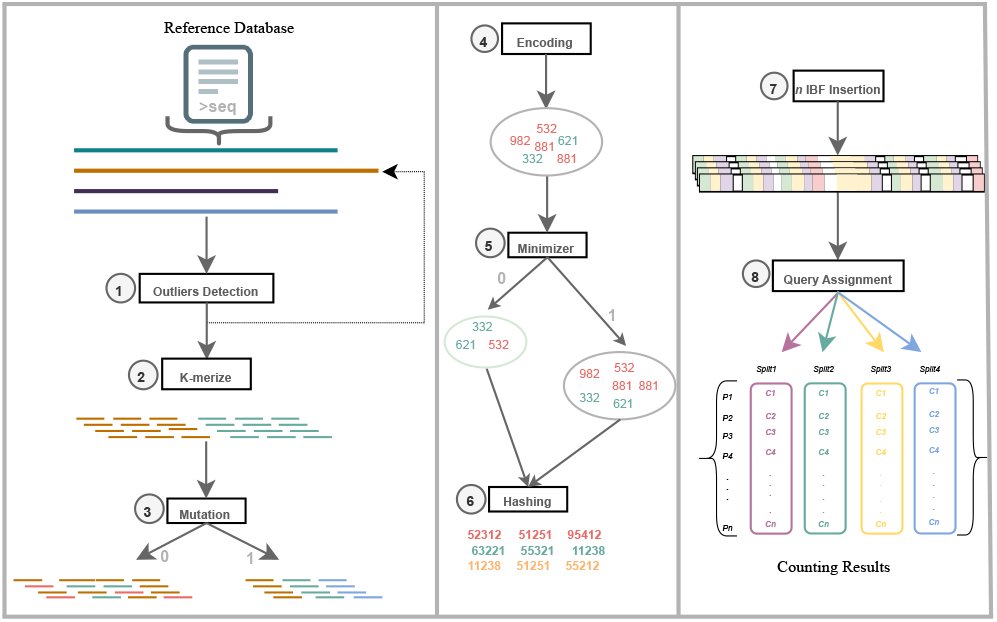
Multi-Indexing Approach Workflow. The user bins are loaded in memory in batches according to the user-defined *M* split size; **(1)** Removing outliers according to the user-defined black list. **(2)** *K*-mer computation of each reference sequence in the loaded split. **(3)** Generating mutation according to user-defined score, if needed (Boolean value 0 or 1). **(4)** Encoding the *k*-mers to the 64-bit integer representation. **(5)** Compute minimizers of each *k*-mers set. **(6)** Hash the encoded values using set of hash functions *H*. **(7)** Insert the *k*-mers into the target IBF; each IBF has the same size as the split size (the last IBF contains the rest of the sequences). **(8)** Query assignment of input peptides against the built index. The query and assignment step is repeated until all IBFs are queried without outliers. The different colors represent the different splits/IBFs.

#### Multi-indexing approach workflow

The workflow begins with the preparation of input datasets, including the reference database and MS/MS spectrum samples, provided in MGF or RAW format (with RAW files converted to MGF). Next, the input protein sequences are *k*-merized to a specified length *k* and inserted into the target IBF index. *De novo* peptides are then sequenced from the input MS/MS spectra using *de novo* sequencing algorithms. These peptides are subsequently *k*-merized at the same length *k* and searched within the correspondingly built index (Suppl. Mat. Figure S2). In the final step, the taxonomy of each sample is determined based on the peptide assignment values to protein sequences, with results scored according to (Equation 6).

### Experiments

We conducted three main experiments to evaluate the performance of our tool in searching and classifying peptide sequences within large protein databases. The results are presented in subsections, which focus on error-free searches and error-tolerant searches that incorporate mutation generation. Finally, we assess potential diagnostic applications, including a case study involving bacterial pathogen detection.

#### Error-free classification

The Mix24X metaproteomics sample [Map+23] contains MS bacterial datasets along with their corresponding references. Initially, these samples were sequenced using Casanovo and SMSNet, and the resulting peptides were pooled and filtered using a cutoff threshold 60%. These peptides were then searched against a combined database that included Mix24X target sequences and the NCBI RefSeqBacterial database (1,728,298 protein sequences) [OLe+15a]. RefSeqBacterial, containing many bacterial protein sequences with a high similarity, was intended to simulate a realistic background for database searches in microbiome experiments. The combined database ultimately comprised around 2 million protein sequences.

For our analysis and comparison with pseudo-ground truth peptides, we utilized the published identifications generated using Mascot Daemon software version 2.5.1 (Matrix Science) [BHC09]. The peptide identifications, as described in the Mix24X publication, were performed with the following parameters: full trypsin specificity, allowance for up to one missed cleavage, static modification of carbamidomethylated cysteine (+57.0215), and variable oxidation of methionine (+15.9949). The mass tolerance was set to 5 ppm for parent ions, with MS/MS tolerances of 0.5 Da and 0.02 Da for the LTQ-Orbitrap XL and Q-Exactive HF instruments, respectively. Peptide matches with a Mascot peptide score corresponding to a p-value below 0.05 were retained. Protein identifications were considered valid if supported by at least two unique peptides. The false-positive rate for protein identification, estimated using a reverse decoy database search under the same parameters, was determined to be below 0.1% [Map+23]. To minimize false positives, a blacklist of overrepresented species was constructed, particularly those with many highly similar sequences from the same proteome. This reduces the number of non-target protein sequences that may otherwise appear frequently simply due to sequence redundancy or similarity. These species, characterized by a large number of proteins and amino acids irrelevant to our analysis, were excluded from the database. Examples include *Serratia* (18,717 proteins; 6,961,050 amino acids) and *Serratia nevei* (11,139 proteins; 4,489,936 amino acids).

MegaPX searches were performed with a *k*-mer size of 5, split size of 1000, and an assignment threshold of 99%, with no minimizers applied. Proteins were concatenated into proteomes using a separator character (*) to prevent overlaps between sequences. This bulk concatenation approach significantly accelerates construction compared to inserting proteins individually into the IBF. In both cases, the IBF avoids inserting duplicate *k*-mers, as each *k*-mer is hashed and the corresponding bit positions are set.

#### Error-tolerant classification

The Lung Squamous Cell Carcinoma (LSCC) study [Sat+21] includes multiple experiments and datasets designed for the proteomics analysis of LSCC. The primary objective of this analysis is to identify human-related proteins. The comprehensive dataset encompasses various MS-based analyses focusing on the phosphoproteome, acetylome, and ubiquitylome. Each sample was analyzed using a Thermo Fisher Scientific Orbitrap Fusion Lumos mass spectrometer equipped with a NanoSpray Flex NG ion source. Specifically, the ubiquitylome dataset, consisting of 30 MS/MS data sets, was analyzed and classified using MegaPX. The analysis of the original study aimed to identify proteins in 30 lung cancer samples. *De novo* peptide sequencing for these samples was conducted using both Casanovo and SMSNet resulting in 608,710 peptide suggestions. Following the *de novo* sequencing, peptides with scores below a 50% threshold were filtered out to ensure data quality. The study includes pseudo-ground truth peptides determined from a database search using MS-GF+ v2017.01.27 [KP14]. Peptide-spectrum matches were identified based on the following database search parameters: a 20 ppm precursor mass tolerance (*-t 20ppm*), semi-tryptic digestion (*-ntt 1*), and a target-decoy approach (*-tda 1*) for FDR estimation. The instrument type was set to Orbitrap (*-inst 1*), and fragmentation mode was specified as HCD for Q-Exactive (*-m 3*) or CID for Orbitrap (*-m 1*). The search allowed up to one isotope error (*-ti 0*,*1)* and considered peptides up to 50 amino acids in length (*-maxLength 50*).

#### Evaluation of diagnostic applications

We searched the Mix24X peptides against the complete protein database of E. *coli* (strain K12) and *Shigella flexneri*, both downloaded from UniProt [Apw+04]. The dataset contains a total of 6,460 proteins in E. *coli* and 63,867 in *Shigella flexneri*. Peptides were searched using a *k*-mer size of 5 and assignment thresholds of 85% and 95%. The search was performed without mutation generation, as gap-based searches were not required. MegaPX utilized a split size of 1,000 user bins per run.

In another experiment, we applied our approach to real bacterial pathogens. In [Kar+20b], species-unique peptide biomarkers for respiratory tract pathogens including *Streptococcus pneumoniae, Haemophilus influenzae, Moraxella catarrhalis*, and *Staphylococcus aureus* were identified. The study highlighted the most promising candidate peptide biomarkers for these four species. In our experiment, we sequenced the corresponding MS datasets (data are available via ProteomeXchange with the identifier PXD014522) using Casanovo and then searched the identified peptide biomarkers within the target 50-reference proteins. This served two purposes: first, to demonstrate that Casanovo can accurately sequence unique peptides in a pathogenic scenario, and second, to show that MegaPX, utilizing the *k*-mer approach (*k=5*), can correctly identify the target reference protein based on these unique peptides. For this experiment, we specifically searched for unique peptides to assess the accuracy of approximate string matching in classifying species-specific unique sequences; we aimed to demonstrate that approximate string matching can reliably classify peptides as species-specific while also being capable of correctly assigning exact matches when no mutations or substitutions are present. This highlights the method’s ability to balance flexibility with precision in sequence classification.

## Results

### Performance evaluation of error-free peptide assignment

All analyses in this section were performed using the Mix24X metaproteomics dataset, which features a known taxonomic composition. This dataset served as a benchmark to evaluate the performance of MegaPX and compare it to alternative methods such as DIAMOND. Both *de novo* peptides (69,656 unique sequences) and Mascot-derived peptides (77,434 unique sequences) were analyzed using a combined reference database (around 2 million sequences). Given that IBF searches are approximate, it is essential to demonstrate the tool’s ability to search within highly similar sequences and accurately assign hits. To further explore this capability, we constructed proteomes from individual proteins, moving from the protein level to the species level. The primary purpose of combining proteins into proteomes was to showcase MegaPX’s capability to accurately assign hits for short peptides within extensive search spaces while utilizing minimal memory usage.

The first search, performed with MegaPX using *de novo* peptides, was completed in 4.68 minutes, consuming 2.18 GB of memory (Suppl. Mat. Table S1). In comparison, the second search, using Mascot peptides required 4.8 minutes and the same memory allocation. Memory usage could potentially be optimized further by reducing split sizes.

We applied a sequence similarity analysis to assess the overlap between *de novo* and Mascot-derived peptides. Only 580 peptides matched exactly between the two methods. However, global alignment [Coc+09] revealed a high degree of similarity, with alignment scores ranging from 70% to 98%. This discrepancy is largely attributed to minor differences in peptide lengths or natural sequence variability introduced by *de novo* sequencing algorithms. To address this, we employed a *k*-mer approach (k=5), which improved overlap detection by identifying partial matches and shared sequence motifs, thus, compensating for the drawbacks of *de novo* sequencing. Figure 3 presents the top 35 taxonomic classifications using MegaPX, successfully assigning *de novo* peptides to the correct strains for the top 23 samples. Strains such as *Pseudomonas putida KT2440* and *Sphingomonas wittichii RW1* dominate the rankings due to their high peptide counts. Conversely,*Staphylococcus_carnosus_subsp_carnosus_TM300* was underrepresented, as its shorter proteins resulted in fewer peptide hits. Next, we compared the counts between Mascot-derived and *de novo* peptides (Suppl. Mat. Figure S3). The results show that *de novo* peptides achieve a lower number of matches to target strains. However, as expected, database-derived peptides demonstrate higher match precision. This finding highlights an additional application of MegaPX as a screening tool. By using MegaPX to assign preliminary hits, researchers can refine their analyses by focusing on high-confidence matches for database searches, reducing the runtime by narrowing subsequent database searches to a subset of relevant references. Nevertheless, MegaPX, when used with*de novo* peptides, still identifies a substantial number of peptide hits.

**Fig. 3.**
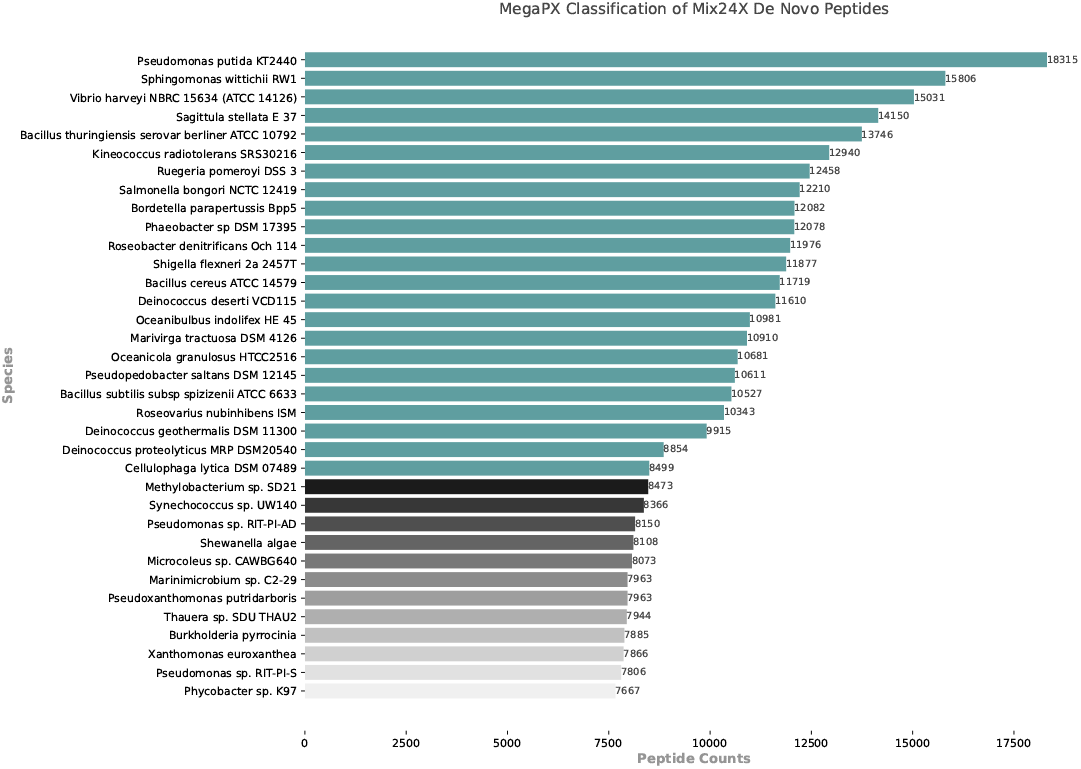
Classification results of *de novo* peptides for the Mix24X sample using MegaPX. The plot shows the top 35 strains ranked by peptide counts. Sequences were queried using *k=5* and an assignment threshold of *D=0*.*99*. MegaPX successfully assigned peptides to the correct strains, with strains like Pseudomonas putida KT2440 and Sphingomonas wittichii RW1 receiving the highest peptide counts.

DIAMOND (version 2.1.9) [BXH14] was tested on the same dataset for comparison. The database was built in 10.3 seconds, and the query search on the pre-built database using *blastp* was completed in 93.21 seconds (Suppl. Mat. Table S1). DIAMOND aligned three queries and four queries when the search was restricted to target proteins. These results highlight the limitations of DIAMOND when handling proteome long sequences with short peptide sequences, emphasizing the advantages of MegaPX’s *k*-mer-based approach in capturing a high number of sequence matches. To address this limitation and allow for a fair comparison - without relying on the approach of concatenating proteins into a single proteome with marker sequences between them - we reconstructed the individual protein sequences from the Mix24X proteomes and performed the search at the protein level rather than the proteome level. We first searched de novo peptides against the Mix24X reference proteins using *DIAMOND*. In this test, the tool reported 6 pairwise alignments, with 4 unique peptide queries successfully aligned. For a more realistic scenario, we performed searches using Mascot-identified peptides (pseudo-ground truht), querying exclusively against target reference proteins with *MegaPX* and *DIAMOND*. This resulted in a total of 178,520 pairwise alignments, with 40,014 peptide queries successfully aligned. Of the 90,616 Mascot peptides, DIAMOND could only align 40,014, representing 44.15% of the complete set of peptides. In contrast, MegaPX demonstrated superior performance, successfully mapping all peptides to their respective references by utilizing the *k*-mer counting technique. Figure 4 illustrates the number of assigned peptides for each target species, showing that MegaPX consistently outperformed DIAMOND across nearly all species. This highlights the advantage of *k*-mer-based approaches in efficiently querying short peptide sequences against large reference databases.

**Fig. 4.**
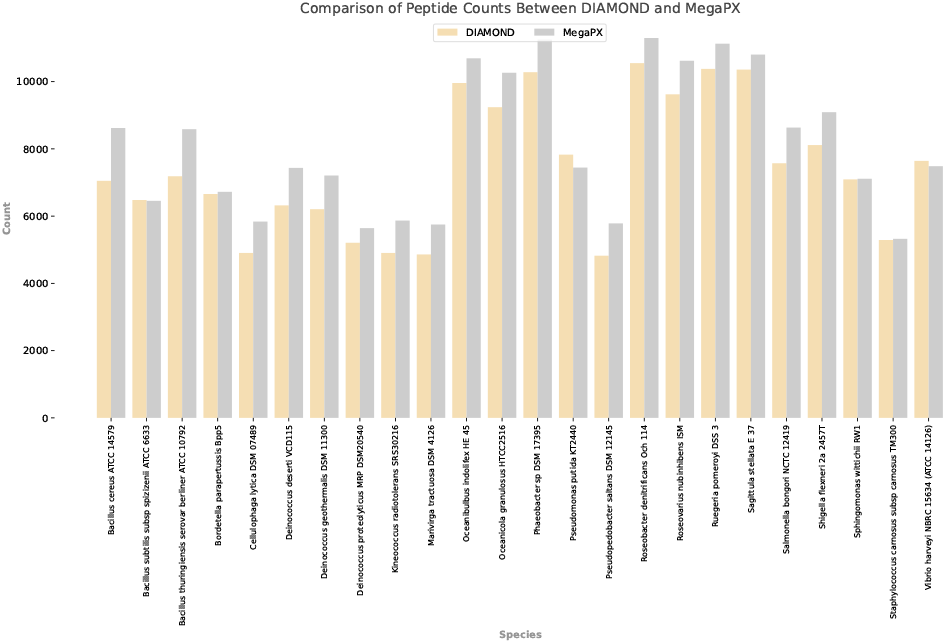
Comparison of peptide counts between DIAMOND and MegaPX. Peptide counts were identified using DIAMOND and MegaPX searches with Mascot’s peptides against Mix24X targets, showcasing the peptide identification capabilities of each method across diverse species.

MegaPX demonstrated its scalability by successfully searching the extensive NCBI_nr database, containing 707 million proteins (339 GB). This search, using a standard notebook with five *k*-mer extraction threads, completed in approximately 18 days while consuming only 0.177 GB of RAM (Suppl. Mat. Table S1). In comparison, DIAMOND faced memory constraints during database building, requiring over 4 GB of RAM and failing to complete the task. MegaPX’s ability to handle large datasets with minimal resources underscores its utility for large-scale metaproteomics studies. The purpose of searching against NCBI_nr was mainly to demonstrate its potential as a screening tool, capable of identifying top-hit sequences. These hits can then be used to narrow down the database for more focused searches or to guide *de novo* sequencing, optimizing both search efficiency and resource usage, this use case could also contribute to the development of advanced algorithm designs for *de novo* sequencing.

### Error-tolerant classification

#### Lung squamous cell carcinoma

To evaluate the ability of MegaPX to identify peptides with high levels of mutations, we conducted a search against a cancer dataset using the mutation mode based on BLOSUM matrices. This dataset contains numerous modifications, making accurate peptide assignment particularly challenging, as mutations can occur at any amino acid position. Notably, the LSCC datasets include pseudo-ground truth peptides identified using the published pipeline, providing an ideal scenario for analyzing shared peptides between SMSNet and Casanovo *de novo* peptides and the pseudo-ground truth peptides. Figure S4 in the Supplement provides a comparison example of shared peptides between pseudo-ground truth and *de novo* sequenced peptides within two LSCC samples. The *de novo* peptides, derived using SMSNet and Casanovo, have been filtered for duplicates and subjected to a score threshold of 50%. In contrast, pseudo-ground truth peptides may include duplicates and are not restricted by a score cutoff. The visualization underscores the overlap and exclusivity of peptides between datasets. Next, the *de novo* collected peptides are searched against the human reference sequence proteome database [OLe+15b]. All references undergo a mutation to a *k*-mer size of 5 with a mutation score of 25 and an assignment threshold of 99%. The original study contains the number and name of the identified proteins for each sample. Figure 5 displays the number of shared proteins identified in each sample. Shared proteins are those identified by MegaPX and are also present in the pseudo-ground truth. MegaPX successfully identified a significant number of proteins from the previous study. However, the pseudo-ground truth results include duplicate proteins and others not present in our reference datasets, as the customized reference database used in the original study was not published. Despite this limitation, MegaPX was able to identify additional proteins from the human proteome reference database. Including these proteins in the results would not allow for a fair comparison with the original publication, as the specific reference database they used was not specified. Therefore, we focused solely on comparing the number of shared peptides/proteins between MegaPX and the published results. Our analysis showed promising results in protein identification scenarios, which facilitates one-to-one identification and peptide assignment of *de novo* peptides with mutation generation. Almost all samples exhibit a high number of matched proteins between cancer and MegaPX results, with a identification recall averaging around 97%, except for four samples (05CPTAC _02, 03CPTAC _01, 03CPTAC _02, 04CPTAC _01) which display protein matching ratios ranging from 57% to 68% (Suppl. Mat. Table S2). These outliers may indicate discrepancies caused by differences in sample quality, experimental conditions, or limitations in the reference datasets. The overall high percentages for most samples demonstrate the robustness of MegaPX in identifying proteins from the original study. The complete search is performed in about 16 minutes with less than 0.5 GB memory consumption with 114 IBFs (Suppl. Mat. Table S1).

**Fig. 5.**
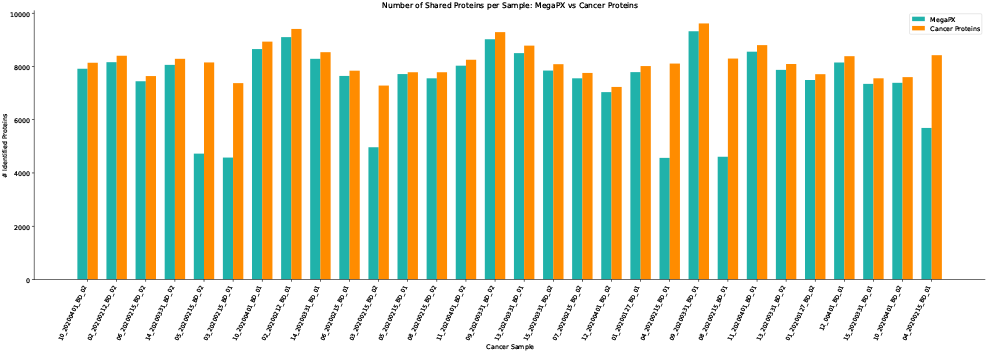
Comparison of the Number of Proteins Identified by MegaPX and the Ground Truth. This figure shows the number of identified proteins per LSCC sample, comparing results from MegaPX with those reported in the original publication based on pseudo-ground truth classifications. To ensure a fair comparison despite differences in reference databases, only shared proteins that were identified by MegaPX and also mentioned in the cancer study were considered.

#### Evaluation of diagnostic applications

##### Sample search against highly similar proteomes

In order to evaluate diagnostic applications, we used the Mix24X dataset, derived from a lab-assembled mixture sample covering 20 genera, including *Shigella flexneri*, a species closely related to *Escherichia coli* with a high degree of proteome sequence overlap. Our goal was to assess the specificity and sensitivity of the diagnostic method in distinguishing between highly similar species. We used MegaPX to search 69,656 *de novo* sequencing-derived peptides from Mix24X against the E. *coli* and *Shigella flexneri* UniProt protein database. The search completed in under one minute, utilizing only 102 MB of memory in both scenarios. The results shown in Figure 6 indicate that the combination of MegaPX and *de novo* peptides proves to be highly effective, as it assigns the highest number of peptides to *Shigella flexneri*. This demonstrates a strong use case for MegaPX, particularly in distinguishing between highly similar proteins with high assignment accuracy. Notably, the tool performs consistently well across both high (95%) and low (85%) assignment threshold scenarios, confirming its robustness and reliability in peptide identification and classification.

##### Species-unique peptide biomarkers

**Fig. 6.**
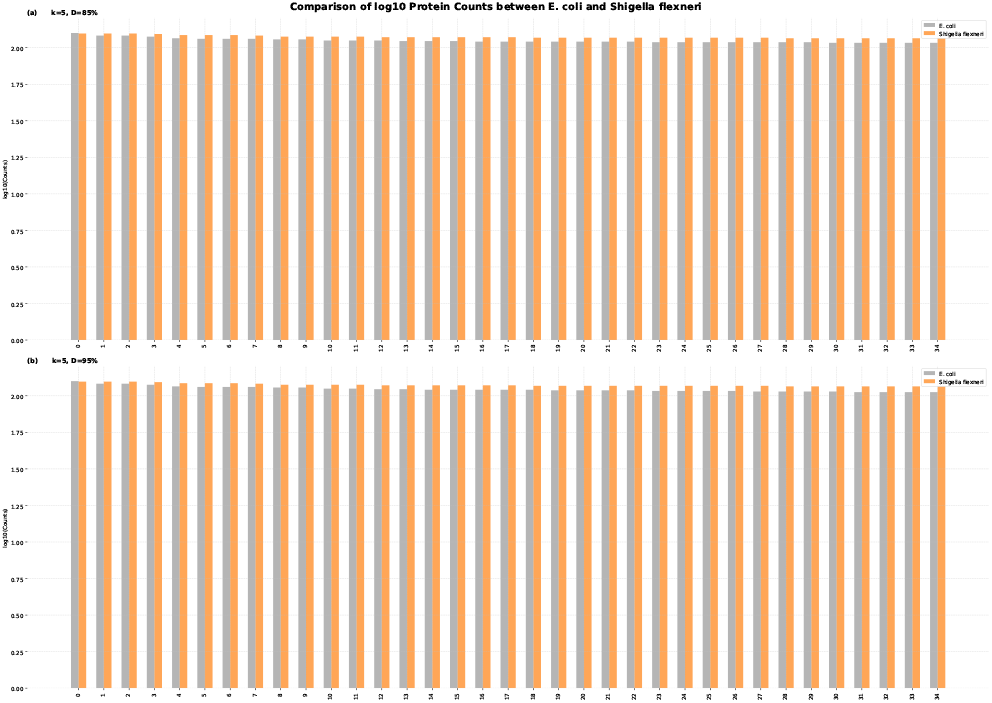
Log10-transformed protein counts comparison between E. *coli* and *Shigella flexneri* under different *k*-mer size and assignment threshold conditions. *k*-mer size = 5, assignment threshold = 85%, (b) *k*-mer size = 5, assignment threshold = 95%. The top 35 proteins are ranked by abundance, with E. *coli* shown in gray and *Shigella flexneri* in orange.

To evaluate Casanovo’s ability to identify species-unique peptide biomarkers in a pathogenic scenario, we applied our approach to datasets of bacterial respiratory pathogens, including *Streptococcus pneumoniae, Haemophilus influenzae, Moraxella catarrhalis*, and *Staphylococcus aureus*. This experiment builds on the work of [Kar+20b], which identified promising candidate peptide biomarkers for these species. Figure 7 presents the number of sequences identified per dataset, along with their corresponding search engine scores from Casanovo. Using Casanovo, we successfully *de novo* sequenced 28 species-unique peptides as exact matches, demonstrating the high precision in peptide identification. Across all datasets, sequence scores were consistently high (*>*80%), ensuring confidence in the obtained results.

**Fig. 7.**
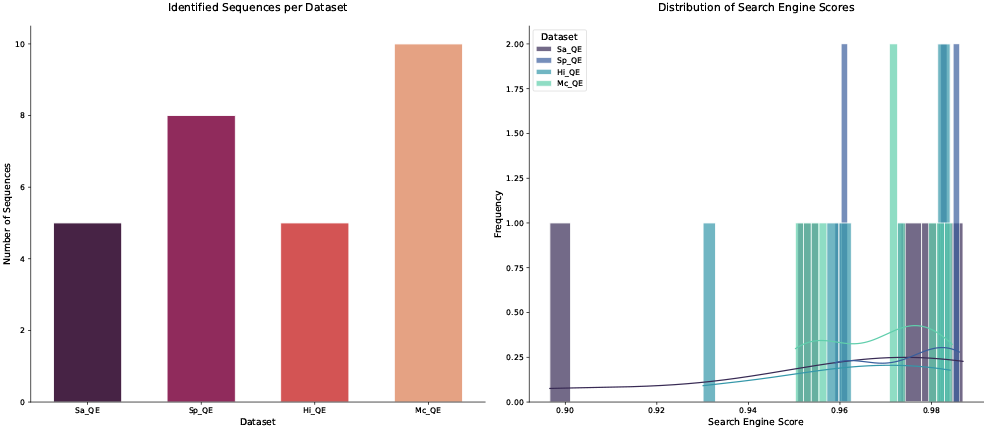
Identified sequences and their corresponding search engine scores across datasets. The bar plot (left) represents the number of sequences identified for each dataset (*Sa_QE, Sp_QE, Hi_QE, Mc_QE*). The histogram (right) shows the distribution of search engine scores, illustrating the frequency of different confidence levels across datasets. *Sa_QE* corresponds to *Staphylococcus aureus, Sp_QE* to *Streptococcus pneumoniae, Hi_QE* to *Haemophilus influenzae*, and *Mc_QE* to *Moraxella catarrhalis*.

The distribution of search engine scores further confirms that the majority of identified peptides were assigned high confidence values. In addition to exact matches, Casanovo identified numerous partial sequence matches to the original peptide biomarkers (e.g., NVATDANHSYTSR and **AN**NVATDANHSYTSR, AVVYNNEGTK and AVVYNNEGTK**VELGGR**). However, we did not consider these partial sequences, as our primary objective was to demonstrate the capability of Casanovo for exact peptide sequencing and to validate the diagnostic applicability of MegaPX for pathogen classification.

To assess *de novo* sequencing accuracy and consistency at the amino acid level, we analyzed amino acid scores across all datasets. Figure 8 illustrates the distribution of these scores across sequence positions and residue types. The results indicate that *de novo* sequencing confidence remains consistently high across different peptide positions, with most residues achieving scores close to 1.0. This suggests that Casanovo reliably assigns amino acids with minimal uncertainty, even in complex bacterial datasets. We further found variations in amino acid confidence across different residue types, providing insights into peptide composition factors that sequencing reliability. These variations may be attributed to amino acid properties or fragmentation patterns. When applying MegaPX searches against the protein reference sequences, all peptides were correctly assigned to their respective target reference sequences (Table 1). In summary, our results demonstrate the reliability of Casanovo in sequencing species-unique peptides while ensuring accurate reference assignment using MegaPX.

**Table 1.**
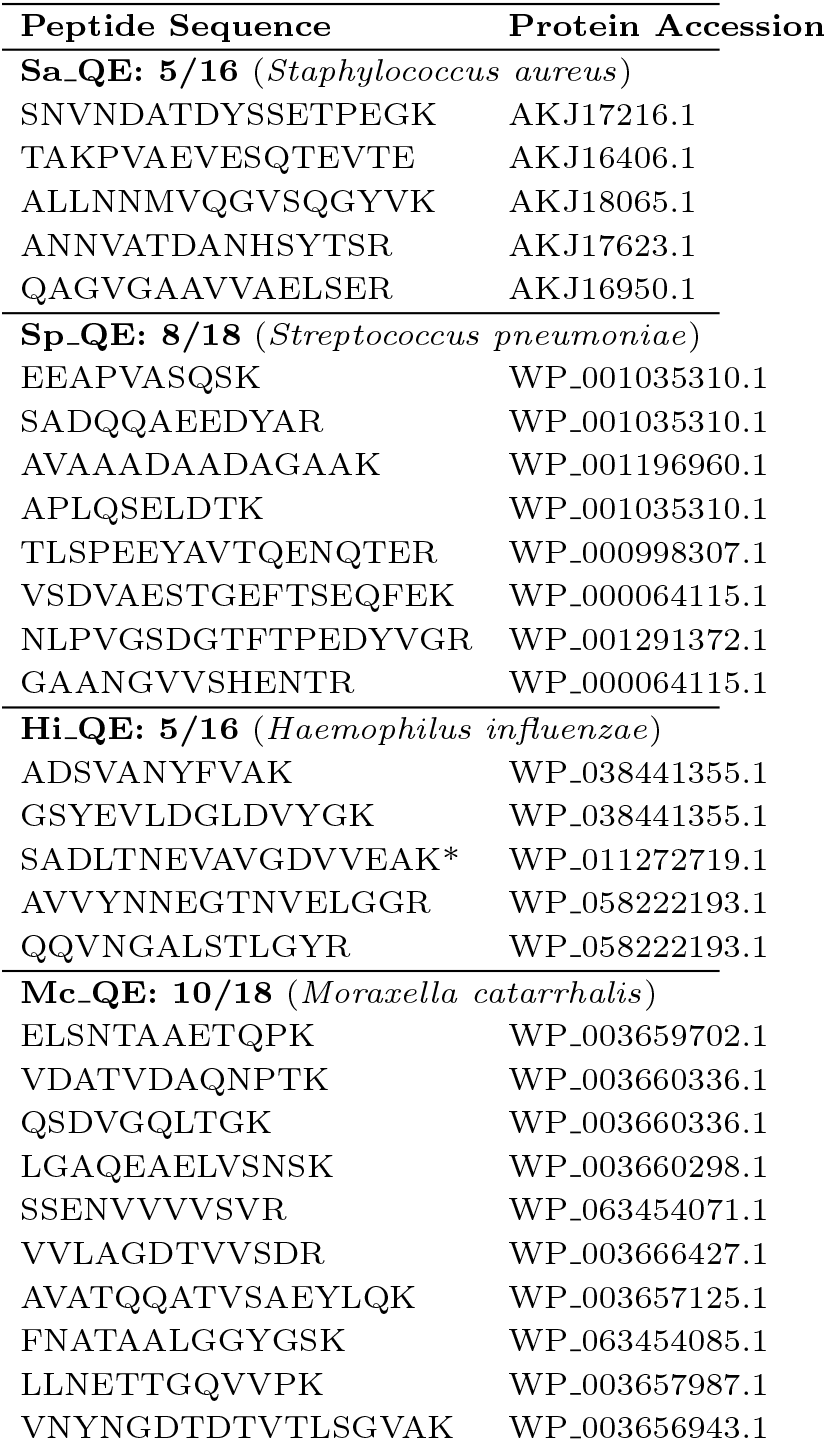
Identified peptide sequences and their corresponding protein accessions across four bacterial pathogens. *Sa_QE* corresponds to *Staphylococcus aureus, Sp_QE* to *Streptococcus pneumoniae, Hi_QE* to *Haemophilus influenzae*, and *Mc_QE* to *Moraxella catarrhalis*. The number of identified peptides per dataset is shown in the format (# identified / # total).

**Fig. 8.**
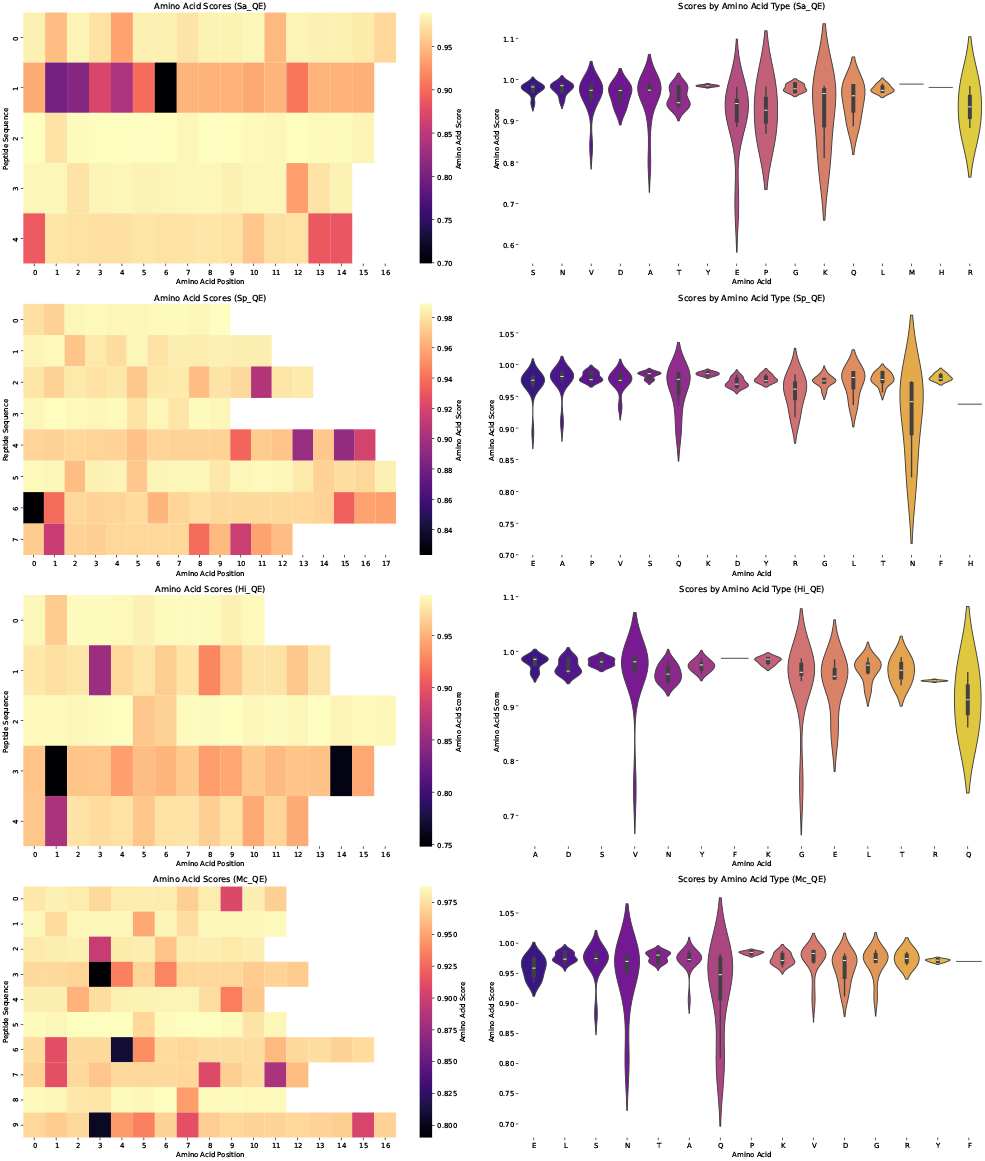
Amino acid scores for identified peptide sequences across datasets. The heatmaps represent amino acid scores at different sequence positions for each dataset (*Sa_QE, Sp_QE, Hi_QE, Mc_QE*), where higher scores indicate greater confidence in sequencing accuracy. The accompanying violin plots display the distribution of amino acid scores grouped by residue type, highlighting sequence composition trends. *Sa_QE* corresponds to *Staphylococcus aureus, Sp_QE* to *Streptococcus pneumoniae, Hi_QE* to *Haemophilus influenzae*, and *Mc_QE* to *Moraxella catarrhalis*.

## Discussion

The development of MegaPX addresses significant challenges in metaproteomics, particularly those related to the taxonomic composition of microbiomes and the limitations of traditional database-dependent peptide identification methods. Our results demonstrate that not only is MegaPX efficient in handling large datasets but also significantly outperforms conventional tools like DIAMOND in both memory usage and hits assignment, particularly in scenarios involving large, complex databases.

One of the key strengths of MegaPX is its ability to perform rapid searches within massive databases, such as the NCBI_nr, which contains over 700 million proteins. The ability to complete these searches in approximately 18 days while consuming only 0.177 GB of RAM highlights the tool’s efficiency and scalability. This contrasts sharply with traditional approaches that often require much more computational resources for similar tasks. Applying this approach on high-performance computing systems would further reduce the runtime by allowing for larger split sizes and increased parallelization. However, this aspect is beyond the scope of our current work, as our primary focus is on enabling accurate classification on standard hardware configurations. Furthermore, the performance of the tool in the Mix24X dataset showcases its ability to accurately assign hits in complex metaproteomic environments. By effectively searching through different databases and correctly identifying short peptides with minimal overlap with traditional database searches, MegaPX demonstrates its robustness and precision.

When applied to the Lung Squamous Cell Carcinoma dataset, MegaPX showed promising results in the identification of human-related proteins, even in the presence of extensive post-translational modifications. Its high average protein matching rate of approximately 97% between the pseudo-ground truth and MegaPX underscores its reliability. Notably, MegaPX’s ability to handle mutation generation and still achieve accurate peptide assignment is particularly beneficial for cancer studies where mutations are frequent and can occur at any amino acid position. the context of diagnostic applications, MegaPX, in combination with *de novo* peptides, effectively assigns the highest number of peptides to *Shigella flexneri*, demonstrating its ability to distinguish highly similar proteins with high assignment accuracy. These findings underscore MegaPX’s potential for precise protein differentiation, even in complex datasets where sequence similarity presents a challenge, making it a valuable tool for high-confidence peptide assignments. The identification of unique peptides across four bacterial species (Table 1) highlights the potential of this approach for pathogen classification using mass spectrometry data. Notably, the high search engine scores (Figure 7) and consistent amino acid confidence levels (Figure 8) confirm the robustness the sequencing and assignment workflow. These findings highlight MegaPX’s capability to support rapid and reliable protein identification, which is essential for developing targeted diagnostic approaches. In general, MegaPX represents a significant advancement in metaproteomics, offering a fast, memory-efficient, and accurate tool for the identification and classification of peptide sequences. Its performance across different datasets and its ability to adapt to various challenges such as large search spaces, high mutation rates, and complex post-translational modifications—underscore its potential as a valuable resource in both research and clinical applications. MegaPX serves as an effective preliminary screening tool for metaproteomics workflows. Its tabular output enables filtering proteins based on peptide identification counts, allowing the generation of smaller, more targeted proteome databases from large-scale databases like NCBI_nr. These refined databases can then be employed in subsequent database search applications, significantly improving both detection accuracy and reducing the runtime of traditional search algorithms. Future optimization of MegaPX could improve processing speed and expand its applicability to other omics fields, such as proteogenomics, which, like metaproteomics, is challenged by a vast search space. Additionally, integrating *de novo* peptide sequencing with MegaPX into a single workflow could provide further benefits to the users.

## Conclusion

The results presented in this work underscore the effectiveness of *k*-mer-based taxonomy classification as a robust method for indexing and querying amino acid sequences, enabling fast and efficient classification. Our study also highlights the promising potential of integrating *de novo* peptide sequencing with classification tasks, offering a viable approach to reducing false positives commonly associated with approximate membership query (AMQ) data structures. Moreover, the successful incorporation of mutated databases into the classification process demonstrates a reliable mechanism for accurately assigning peptides to target proteins or reference sequences, even in the presence of sequencing errors or mutations related to viruses or cancer. This integration is crucial for maintaining accuracy in challenging metaproteomics analyses. Our efforts to optimize this process for fast and memory-efficient execution have shown that AMQ data structures offer a robust solution that balances accuracy with efficient runtime performance. These findings pave the way for the development of improved methods in bioinformatics, meeting the growing demands of modern genomic and proteomic research.

## Acknowledgements

The authors thank the SeqAn team (Freie Universität Berlin) for providing the C++ library and for the valuable discussions and comments on AMQ data structures.

## Notes

### Competing Interest Statement

The authors have declared no competing interest.

### Funding

- German Research Foundation (DFG): MU 4430//2-1

