## Supplementary Material for "MegaPX: fast and space-efficient peptide assignment method using IBF-based multi-indexing"

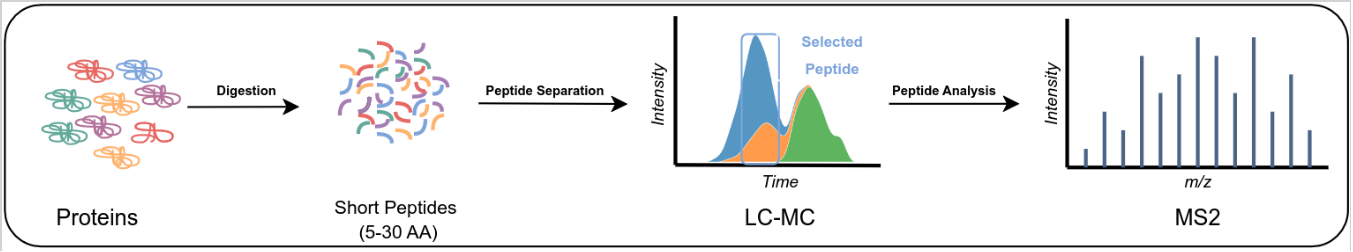


**Figure S1:** Bottom-Up Proteomics Workflow. Target proteins are digested by trypsin to cleave them into short peptides of length 5 to 30 AA. These peptides are then separated by LC and subsequently analyzed with the mass analyzer. The final MS/MS step describes the spectra generated for computer-aided analysis.

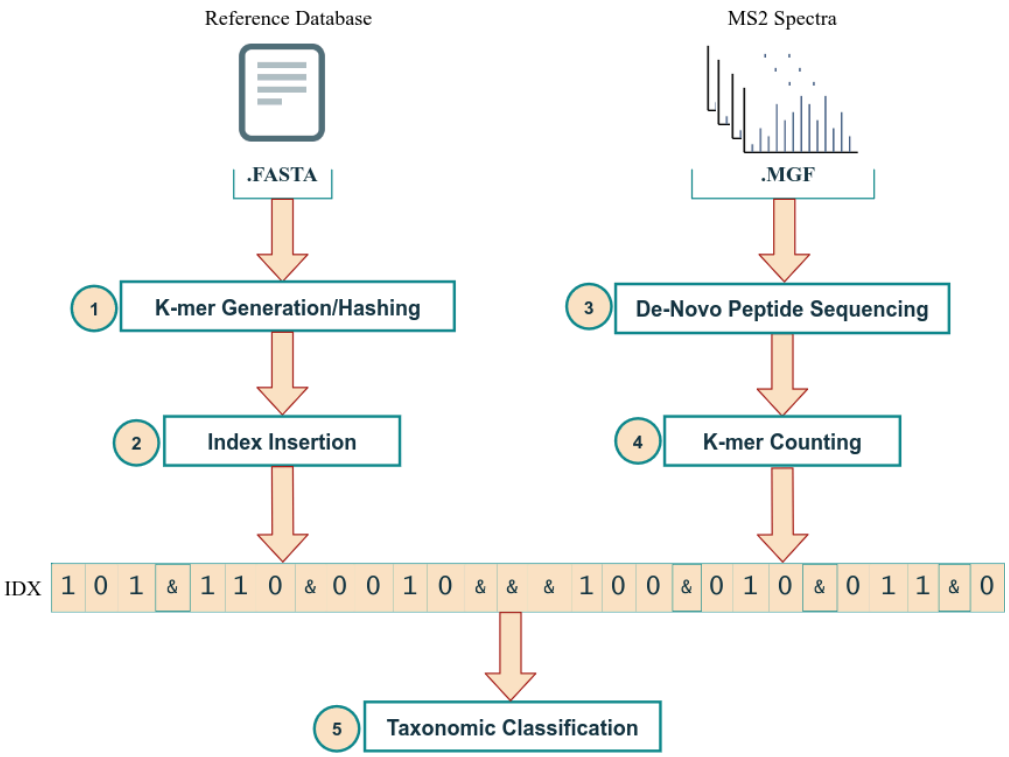


**Figure S2:** Classification Workflow. **(1)** Generate *k*-mers from each reference sequence and apply arithmetic coding with hashing to the *k*-mers. **(2)** Insert all generated *k*-mers into the corresponding index the user selects. **(3)** *de novo* peptide sequencing of the input spectra samples. **(4)** Counting the number of *k*-mers in the generated index. **(5)** Assign the counts and create a report on the classification of the last output of the taxonomy.

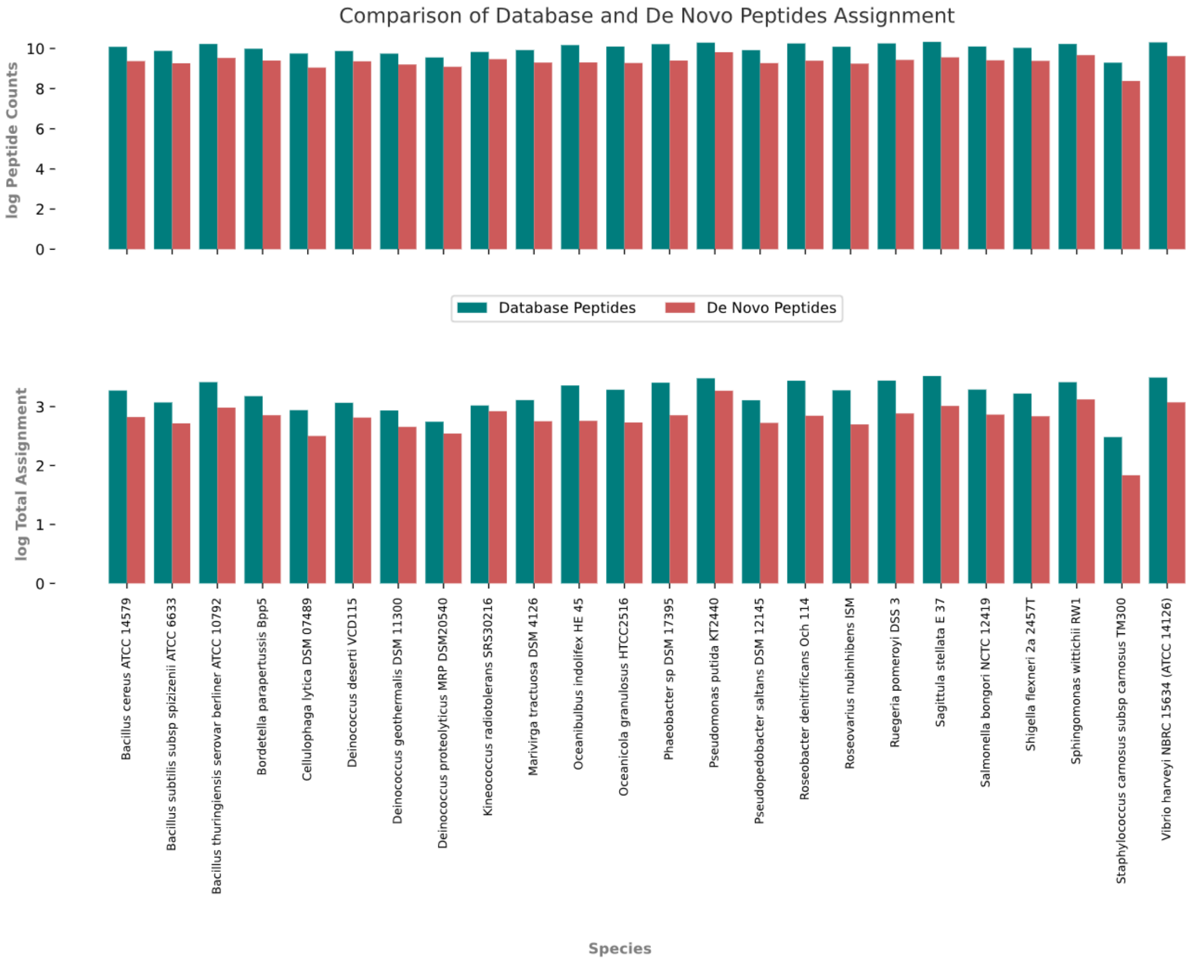


**Figure S3:** Comparison of species matches between *de novo* and database peptides for the Mix24X sample. The top plot displays the peptide counts on a logarithmic scale, showing that *de novo* peptides yield a high number of matches to target strains, likely due to the larger pool of *de novo* peptide sequences.The bottom plot shows the total number of species assignments per peptide set on a logarithmic scale. Since individual peptides may match multiple species, values greater than the total peptide count can occur. This metric reflects the overall assignment coverage and redundancy, not precision.The analysis was conducted with *k=5* and an assignment threshold of *D=0.99*, excluding minimizer computation.

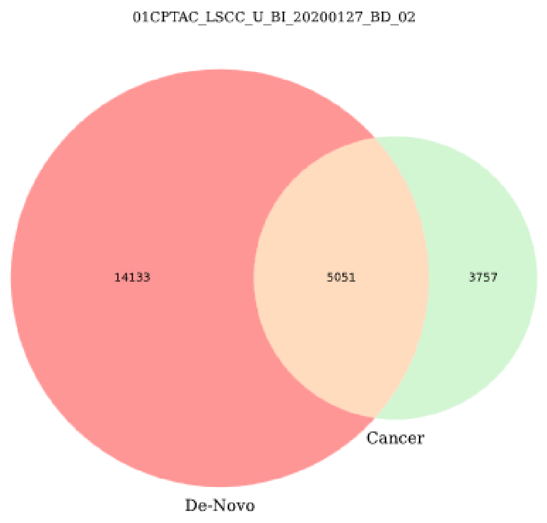


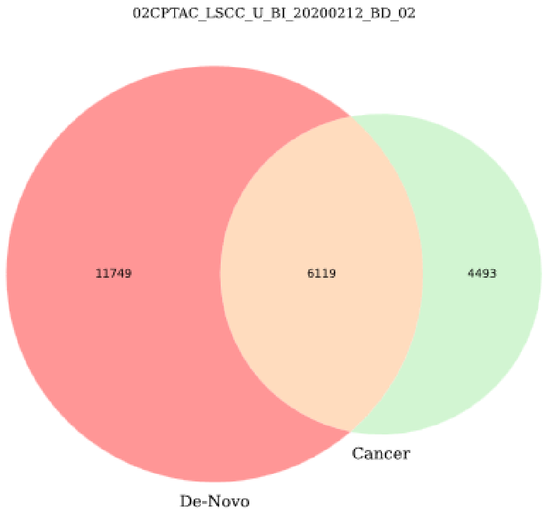


**Figure S4:** Set Membership of Shared Peptides in LSCC samples 01CPTAC_2 and 02CPTAC_2. This figure illustrates the overlap of shared peptides between *de novo* peptides and pseudo-ground truth peptides in LSCC samples 01CPTAC_2 and 02CPTAC_2, where *de novo* peptides have been filtered for duplications and a score threshold of 50%.

| Experiment | Peptides | Reference DB | Run Time | Memory (GB) |
|---|---|---|---|---|
| Mix24X-MegaPX | *de novo* | combined DB | 4.68 min | 2.18 |
| Mix24X-MegaPX | database | combined DB | 4.1 min | 2.18 |
| Mix24X-DIAMOND | database | combined DB | 1.7 min | 2 |
| Mix24X-MegaPX | *de novo* | NCBI_nr | 18 days | 0.177 |
| Mix24X-DIAMOND | *de novo* | NCBI_nr | ——— | $> 4$ |
| LSCC-MegaPX | *de novo* | Mutated Human Proteome | 16.5 min | 0.46 |
| E.coli-MegaPX | *Mix24X de novo* | E.coli proteins | 3.4 sec | 0.102 |

**Table S1:** Runtime and Memory Usage Analysis for Different Experiments. *Combined DB*: RefSeqBacterial proteins combined with the Mix24X targets.

| Sample | MegaPX Hits | Ground Truth | Identification Ratio (%) |
|---|---|---|---|
| 10CPTAC_LSCC_U_BI_20200401_BD_02 | 7914 | 8139 | 97.24 |
| 02CPTAC_LSCC_U_BI_20200212_BD_02 | 8160 | 8402 | 97.12 |
| 06CPTAC_LSCC_U_BI_20200215_BD_02 | 7445 | 7639 | 97.46 |
| 14CPTAC_LSCC_U_BI_20200331_BD_02 | 8060 | 8288 | 97.25 |
| 05CPTAC_LSCC_U_BI_20200215_BD_02 | 4726 | 8150 | 57.99 |
| 03CPTAC_LSCC_U_BI_20200215_BD_01 | 4578 | 7373 | 62.09 |
| 10CPTAC_LSCC_U_BI_20200401_BD_01 | 8653 | 8932 | 96.88 |
| 02CPTAC_LSCC_U_BI_20200212_BD_01 | 9102 | 9409 | 96.74 |
| 14CPTAC_LSCC_U_BI_20200331_BD_01 | 8287 | 8537 | 97.07 |
| 06CPTAC_LSCC_U_BI_20200215_BD_01 | 7644 | 7841 | 97.49 |
| 03CPTAC_LSCC_U_BI_20200215_BD_02 | 4962 | 7283 | 68.13 |
| 05CPTAC_LSCC_U_BI_20200215_BD_01 | 7714 | 7928 | 97.3 |
| 08CPTAC_LSCC_U_BI_20200215_BD_02 | 7554 | 7782 | 97.07 |
| 11CPTAC_LSCC_U_BI_20200401_BD_02 | 8029 | 8252 | 97.3 |
| 09CPTAC_LSCC_U_BI_20200331_BD_02 | 9022 | 9290 | 97.12 |
| 13CPTAC_LSCC_U_BI_20200331_BD_01 | 8502 | 8782 | 96.81 |
| 15CPTAC_LSCC_U_BI_20200331_BD_02 | 7847 | 8086 | 97.04 |
| 07CPTAC_LSCC_U_BI_20200215_BD_02 | 7555 | 7757 | 97.4 |
| 12CPTAC_LSCC_U_BI_20200401_BD_02 | 7037 | 7231 | 97.32 |
| 01CPTAC_LSCC_U_BI_20200127_BD_01 | 7787 | 8016 | 97.14 |
| 04CPTAC_LSCC_U_BI_20200215_BD_02 | 4566 | 8109 | 56.31 |
| 09CPTAC_LSCC_U_BI_20200331_BD_01 | 9322 | 9616 | 96.94 |
| 08CPTAC_LSCC_U_BI_20200215_BD_01 | 4609 | 8296 | 55.56 |
| 11CPTAC_LSCC_U_BI_20200401_BD_01 | 8556 | 8801 | 97.22 |
| 07CPTAC_LSCC_U_BI_20200215_BD_01 | 7872 | 8093 | 97.27 |
| 15CPTAC_LSCC_U_BI_20200331_BD_01 | 8149 | 8387 | 97.16 |
| 13CPTAC_LSCC_U_BI_20200331_BD_02 | 7347 | 7554 | 97.26 |
| 01CPTAC_LSCC_U_BI_20200127_BD_02 | 7385 | 7599 | 97.18 |
| 12CPTAC_LSCC_U_BI_20200401_BD_01 | 7492 | 7710 | 97.17 |
| 04CPTAC_LSCC_U_BI_20200215_BD_01 | 5690 | 8424 | 67.55 |

**Table S2:** Protein Matching Results Between MegaPX and Pseudo-Ground Truth Cancer Proteins. This table summarizes the number of proteins identified by MegaPX in each LSCC sample compared to the pseudo-ground truth cancer proteins. The matching ratio (%) represents the **recall**, calculated as the proportion of pseudo-ground truth proteins that were successfully identified by MegaPX. Specifically, it is computed by dividing the number of overlapping proteins by the total number of pseudo-ground truth proteins. While most samples achieve a high matching ratio (approximately 97%), a few outliers with lower ratios (57–68%) are noted.

**Algorithm 1** Multi-indexing

**Require:** User-defined split size $M$
**Ensure:** All input peptides are queried against the built index
**for** each batch of user bins loaded in memory **do**
    Remove outliers according to the user-defined blacklist
    Compute $k$-mers of each reference sequence in the loaded split
    **if** error-prone search (Boolean value 0 or 1) **then**
        Generate mutation according to user-defined score
    **end if**
    Encode the $k$-mers to the 64-bit integer representation
    Compute minimizers of each $k$-mer set (Boolean value 0 or 1)
    Hash the encoded values using a set of hash functions $H$
    Insert the $k$-mers into the target IBF; each IBF has the same size as the split size
**end for**
Assign queries of input peptides against the built index
Repeat query and assignment step until all IBFs are queried

**Algorithm S1:** Multi-indexing Pseudo-code.
